# Transcriptomic profiling of the human habenula reveals a shared molecular architecture across mood disorders

**DOI:** 10.64898/2026.06.01.728671

**Authors:** Sarah Cameron, Marnie L. Maddock, Samara J. Walpole, Katrina Weston-Green, Kelly A. Newell

## Abstract

The habenula is a critical regulator of monoaminergic and reward circuitry and is increasingly implicated in the neurobiology of mood disorders. Preclinical studies demonstrate that habenula hyperactivity drives depressive-like behaviours and can be reversed by interventions such as ketamine and deep brain stimulation. However, the molecular architecture of the human habenula remains largely unexplored. Here, we applied a transdiagnostic framework to characterise shared and disorder-specific transcriptomic alterations across affective illness. Bulk RNA sequencing was performed on postmortem habenula-enriched tissue from controls (n = 6), major depressive disorder (MDD; n = 6), and bipolar disorder (BPD; n = 6) cases from the Netherlands Brain Bank. Differential gene expression and differential transcript usage (DTU) analyses identified diagnosis- and sex-associated transcriptional changes. Individual diagnostic comparisons revealed modest numbers of differentially expressed genes (MDD: 60; BPD: 66; FDR < 0.05), consistent with limited power and/or subtle disorder-specific effects. In contrast, transdiagnostic analysis combining affective cases (MDD + BPD; n = 12) identified 378 differentially expressed genes, indicating a robust shared molecular signature across mood disorders. Upregulated genes were enriched for potassium channel activity, calcium homeostasis, and Wnt signalling, consistent with altered neuronal excitability, while downregulated genes were enriched for metal ion binding. DTU analysis identified 49 isoform switches, highlighting isoform-specific regulation not captured at the gene level. Biological sex contributed substantially to transcriptomic variation, with 67 differentially expressed genes and 18 isoform switches differing between males and females, including sex-dependent regulation of GPR151, NLGN3, and KIF17, genes known to influence neuronal excitability. Together, these findings support a shared molecular architecture across mood disorders and underscore the importance of transdiagnostic and sex-informed approaches.

## 1. Introduction

The habenula is a small, evolutionarily conserved epithalamic structure that has emerged as a critical regulator of mood, with growing evidence linking its dysfunction to psychiatric conditions (1). It is comprised of medial (MHb) and lateral (LHb) subdivisions that are anatomically and functionally distinct (2). Often referred to as the brain’s “anti-reward” or “disappointment centre”, the LHb exerts top-down inhibitory control over dopaminergic, serotonergic and noradrenergic systems, making it a key point of convergence modulating monoamine transmission (3). The LHb connection to several brain regions involved in emotional regulation, stress (4), motivation (5), sleep (6) and the encoding of negative stimuli (7) suggest it may act as a central driver influencing core neurobiological processes disrupted in depression. The comparatively lesser known MHb has been linked to the regulation of fear and anxiety (8) and has been heavily implicated in nicotine addiction (9). While emerging preclinical evidence suggests a potential role for the MHb in depressive-like behaviours, investigations of this subdivision remain limited, largely due to its small size and relative inaccessibility (10,11).

Preclinical studies provide compelling evidence for a causal role of LHb dysfunction in depression. Remarkably, direct stimulation of the LHb in otherwise healthy rodents is sufficient to induce a depressive-like phenotype, suggesting LHb hyperactivity may be a primary pathology in depression (12,13). Consistent with this, preclinical models of depression demonstrate elevated baseline LHb activity which is reversible by novel therapeutics such as ketamine (14) and by deep brain stimulation (15). At the molecular level, habenula hyperactivity in preclinical models of depression has largely been attributed to disruption of excitatory/inhibitory balance, with enhanced glutamatergic signalling (16,17), altered calcium-dependent signalling (18), and maladaptive astrocyte functioning (19) identified as key contributors. Despite consistent preclinical findings, the molecular underpinnings of habenula dysfunction in human depression remain poorly defined. Notably, the morphological organisation of the human habenula differs from that of rodents, suggesting species-specific molecular and functional distinctions (20). Additionally, in humans, the LHb comprises a substantially larger proportion of the whole habenula complex (∼95%) than in rodents (∼60%) (20,21). The relative size difference may indicate a more complex modulation within the human LHb, emphasising the need for clinical research.

Clinical studies report altered habenula activation, connectivity and volume in individuals with depression (22–24). However, these findings are largely derived from neuroimaging methods that are constrained by the small size of the structure and offer limited insight into the molecular landscape of the human habenula. Notably, habenula dysfunction is also clinically relevant to bipolar disorder, in which depressive symptoms dominate the longitudinal course of illness and contribute substantially to morbidity and suicide risk (25,26). Importantly, major depressive disorder and bipolar disorder share substantial clinical, genetic, and neurobiological overlap, particularly in relation to depressive symptomatology (27,28). This has led to increasing recognition of mood disorders as existing along a transdiagnostic spectrum, with shared circuit-level dysfunctions that may transcend categorical diagnoses (27,28). In this context, the habenula represents a compelling candidate structure mediating common pathophysiological processes across affective illness. Despite this overlap, bipolar disorder has received comparatively less focused investigation in the context of habenula dysfunction. Early high-resolution MRI work reported reduced habenula volume in unmedicated individuals with bipolar disorder experiencing depressive episodes (29). Although subsequent larger-scale imaging studies failed to detect robust diagnostic group differences in habenula volume but identified associations between right habenula variability and suicidality in bipolar disorder (30). Together, these findings suggest that habenula alterations in bipolar disorder may be state-dependent, clinically heterogeneous, or obscured by methodological limitations inherent to imaging the habenula. Beyond structural imaging, direct functional evidence in humans remains limited; however, invasive electrophysiological recordings from the habenula in individuals with treatment-resistant unipolar or bipolar depression undergoing deep brain stimulation revealed abnormal excitation–inhibition signatures that correlated with depressive symptom severity, providing convergent support for habenula hyperactivity across mood disorders (31).

To date, only a small number of postmortem molecular studies have examined the human habenula, with most being derived from male-only cohorts, leaving it unclear whether habenula molecular profiles differ by sex or carry distinct clinical implications (32,33). Notably, the recent study by Yalcinbas et al. provided the first comprehensive transcriptomic characterisation of the human habenula in schizophrenia, identifying molecularly distinct LHb and MHb cell populations and demonstrating that schizophrenia-associated gene expression changes in the male habenula are largely unique compared to other brain areas. While these findings establish the habenula as a transcriptionally distinct and disease relevant structure in the human brain, they also highlight the broader lack of molecular data in females and across other psychiatric conditions including major depressive disorder and bipolar disorder. This gap is particularly important considering preclinical evidence demonstrating pronounced sex differences in afferent excitatory connectivity to LHb (34). Clinically, women represent two-thirds of the individuals diagnosed with depression (35), and while bipolar disorder shows comparable prevalence across sexes, females more frequently present with bipolar II disorder, which is characterise by predominantly depressive episodes (36).

Together, these findings position the habenula as a critical, yet insufficiently characterised structure involved in the neurobiology of mood disorders. While preclinical studies have delineated robust mechanisms of habenula dysfunction, their translation to the human brain, particularly at the molecular level, remains limited. A deeper understanding of transcriptomic alterations within the human habenula, and how these vary across sex and psychiatric diagnosis, is essential to bridge this translational gap and inform the development of more precise, mechanistic based therapeutics. In this study, bulk RNA sequencing was used to characterise transcriptional alterations in the human habenula across mood disorders. We examined both diagnosis-specific and shared (transdiagnostic) expression patterns across major depressive disorder and bipolar disorder to identify molecular signatures that converge across affective illness. In parallel, we assessed sex-associated transcriptional differences to address the marked sex bias in mood disorder prevalence and the paucity of female representation in existing postmortem studies.

## 2. Methods

### 2.1 Postmortem brain tissue

Postmortem human habenula-enriched tissue was obtained from the Netherland Brain Bank (NBB). Given the high clinical overlap between major depressive disorder and bipolar disorder, the study was designed to enable both diagnosis-specific and transdiagnostic analyses of mood pathology. The cohort consisted of tissue samples from control (n=6), bipolar disorder (BPD) (n=6) and major depressive disorder (MDD) (n=6) cases. Comprehensive demographic and clinical records for all donors was provided by the Netherlands Brain Bank (**Table 1**).

**Table 1.**
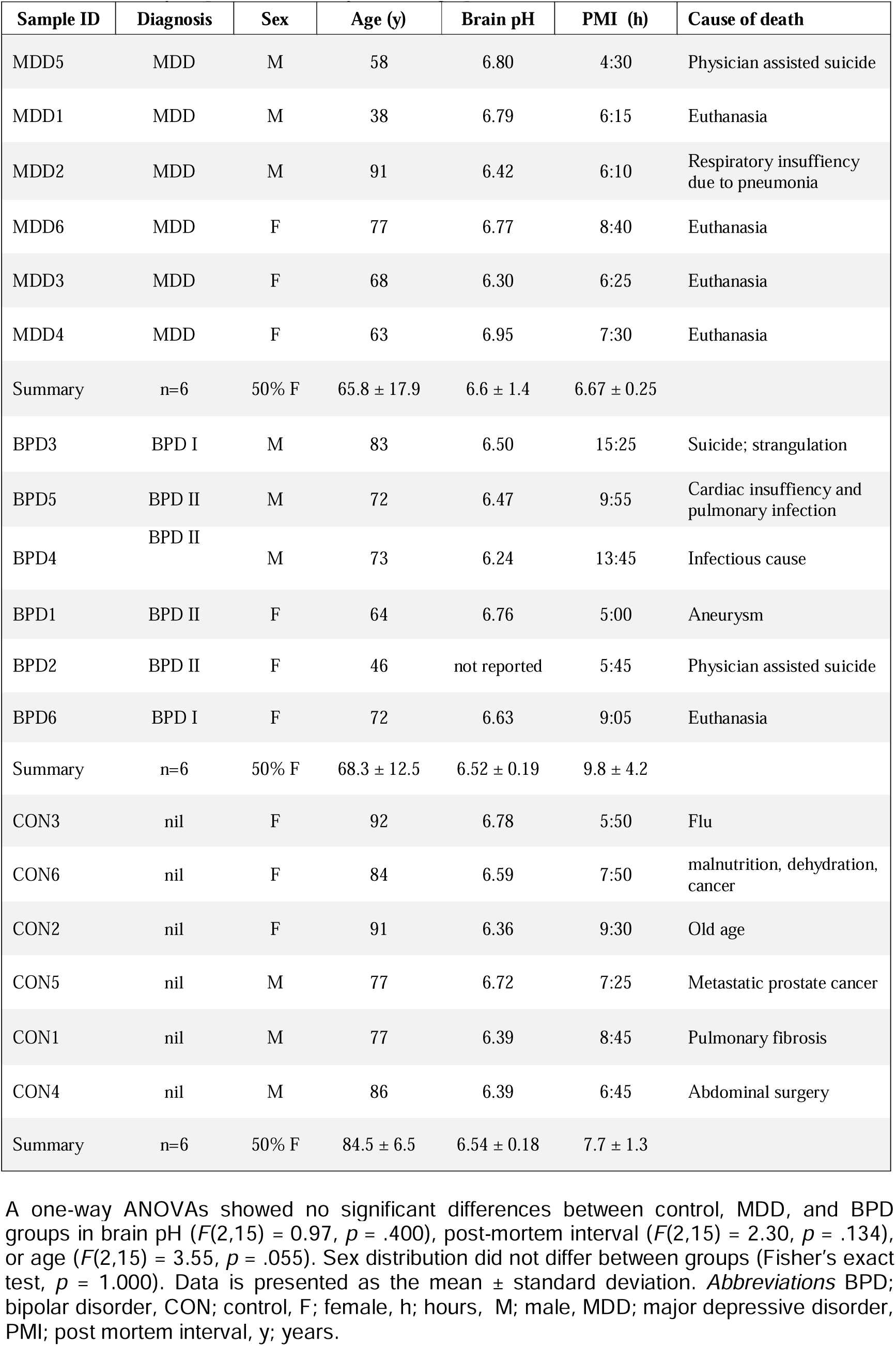
Summary of postmortem subject demographics. A one-way ANOVAs showed no significant differences between control, MDD, and BPD groups in brain pH (*F*(2,15) = 0.97, *p* = .400), post-mortem interval (*F*(2,15) = 2.30, *p* = .134), or age (*F*(2,15) = 3.55, *p* = .055). Sex distribution did not differ between groups (Fisher’s exact test, *p* = 1.000). Data is presented as the mean ± standard deviation. *Abbreviations* BPD; bipolar disorder, CON; control, F; female, h; hours, M; male, MDD; major depressive disorder, PMI; post mortem interval, y; years.

| Sample ID | Diagnosis | Sex | Age (y) | Brain pH | PMI (h) | Cause of death |
| --- | --- | --- | --- | --- | --- | --- |
| MDD5 | MDD | M | 58 | 6.80 | 4:30 | Physician assisted suicide |
| MDD1 | MDD | M | 38 | 6.79 | 6:15 | Euthanasia |
| MDD2 | MDD | M | 91 | 6.42 | 6:10 | Respiratory insufficiency due to pneumonia |
| MDD6 | MDD | F | 77 | 6.77 | 8:40 | Euthanasia |
| MDD3 | MDD | F | 68 | 6.30 | 6:25 | Euthanasia |
| MDD4 | MDD | F | 63 | 6.95 | 7:30 | Euthanasia |
| Summary | n=6 | 50% F | 65.8 ± 17.9 | 6.6 ± 1.4 | 6.67 ± 0.25 |  |
| BPD3 | BPD I | M | 83 | 6.50 | 15:25 | Suicide; strangulation |
| BPD5 | BPD II | M | 72 | 6.47 | 9:55 | Cardiac insufficiency and pulmonary infection |
| BPD4 | BPD II | M | 73 | 6.24 | 13:45 | Infectious cause |
| BPD1 | BPD II | F | 64 | 6.76 | 5:00 | Aneurysm |
| BPD2 | BPD II | F | 46 | not reported | 5:45 | Physician assisted suicide |
| BPD6 | BPD I | F | 72 | 6.63 | 9:05 | Euthanasia |
| Summary | n=6 | 50% F | 68.3 ± 12.5 | 6.52 ± 0.19 | 9.8 ± 4.2 |  |
| CON3 | nil | F | 92 | 6.78 | 5:50 | Flu |
| CON6 | nil | F | 84 | 6.59 | 7:50 | malnutrition, dehydration, cancer |
| CON2 | nil | F | 91 | 6.36 | 9:30 | Old age |
| CON5 | nil | M | 77 | 6.72 | 7:25 | Metastatic prostate cancer |
| CON1 | nil | M | 77 | 6.39 | 8:45 | Pulmonary fibrosis |
| CON4 | nil | M | 86 | 6.39 | 6:45 | Abdominal surgery |
| Summary | n=6 | 50% F | 84.5 ± 6.5 | 6.54 ± 0.18 | 7.7 ± 1.3 |  |

Cases were matched for key demographic variables including postmortem interval, age, brain pH and sex. Psychiatric diagnoses were established prior to death by a medical doctor according to the Diagnostic and Statistical Manual of Mental Disorders (DSM). Any documented lifetime history of psychiatric medication use was obtained from clinical records provided by the NBB. Among affective disorder cases (MDD and BPD), all individuals had documented exposure to antidepressants, including selective serotonin reuptake inhibitors (SSRIs), serotonin norepinephrine reuptake inhibitors (SNRIs), tricyclic and tetracyclic antidepressants, monoamine oxidase inhibitors (MAOIs), and atypical antidepressants. Antipsychotic exposure was present in 58% of cases (quetiapine, olanzapine, lurasidone, and risperidone), while 50% of cases had a history of mood stabiliser use (lithium, carbamazepine, lamotrigine, and sodium valproate).

The study was approved by the University of Wollongong Human Research Ethics Committee (HE2013/069). Clinical information indicated that most BPD cases were diagnosed with BPD II and predominately experienced depressive episodes, with limited history of mania. Postmortem dissection of the habenula-enriched tissue was performed by the Netherlands Brain Bank following their standard protocol (**Figure 1**). Briefly, the pineal gland was retracted with tweezers to visualise the habenula commissure. Approximately 1 cm of tissue surrounding the habenula commissure was dissected antero-laterally to ensure enrichment of the habenula region. Consistent with successful regional enrichment, transcriptomic analysis detected robust expression of established habenula-enriched marker genes including *GRP151, POU4F1, CHRNA3* and *CHRNB3*, which have previously been reported as characteristic markers of the human habenula (20,33,37).

**Figure 1.**
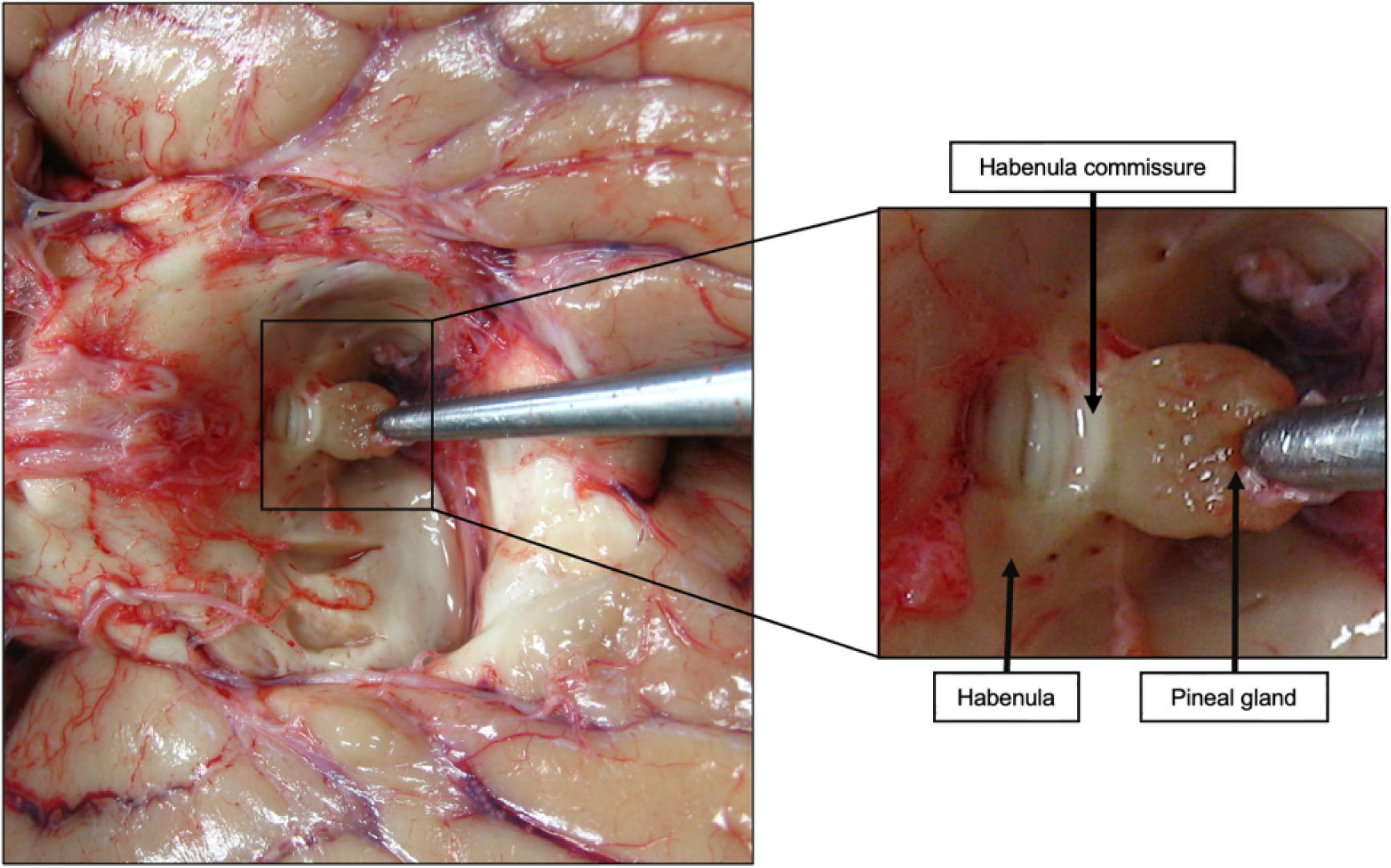
Dissection of the human habenula from postmortem brain tissue. The brain is positioned upside down (resting on the cerebral cortex) with the brainstem and cerebellum flipped anteriorly to expose the pineal gland (left). The enlarged view (right) highlights the anatomical landmarks of the habenula region, including the pineal gland, habenula commissure, and left habenula.

### 2.3 RNA extraction, library preparation and RNA sequencing

Total RNA was extracted from habenula-enriched tissue using the Thermo Fisher PureLink™ RNA Mini Kit (Thermo Fisher Scientific, USA) according to the manufacturer’s protocol. RNA concentration and purity were quantified using a ND-1000 Spectrophotometer (Nanodrop Technologies, Wilmington, DE, USA). Libraries were prepared using the Illumina Stranded Total RNA Prep with RiboZero Plus and paired-end sequencing of 100 bp read lengths was conducted by the Ramaciotti Centre for Genomics (UNSW Sydney, Australia) using the NovaSeq 6000 platform (Illumina inc.). One sample (CON5) had a low RNA integrity number (RIN), indicating low quality RNA and was removed from subsequent analysis.

Raw sequencing reads were processed using the nf-core RNAseq pipeline (version 3.18.0) (38). Read quality was assessed using FastQC, and adapter and quality trimming were performed using Trim Galore. Reads were then aligned to the human reference genome (GRCh38.113) using the STAR aligner. An average sequencing depth of 94 million total reads was obtained per sample, with 97.6% of reads aligning to the genome and 92.2% mapping uniquely. The average sequencing depth did not differ between groups (F(2,3)= 1.35, *p* = .29). Transcript quantification was performed using Salmon, generating gene-level abundance expressed as transcripts per million (TPM). Read alignment metrics and further quality assessment were conducted using MultiQC.

### 2.4 Data Analysis

To examine both diagnosis-specific and shared molecular signatures across affective illness, analyses were conducted at the level of individual diagnoses (MDD, BPD) and using a pre-defined transdiagnostic mood cohort (MDD + BPD), based on established clinical and neurobiological overlap between these disorders (27,28). To compare differentially expressed genes (DEGs) between CON, BPD, MDD and the combined mood group, raw count data was imported into R (version 4.4.2) and analysed using the DESeq2 package (version 1.48.2) (39). Genes with very low expression (less than 10 counts) were excluded from further analysis to minimise noise and false positives, resulting in 38,872 genes retained for analysis. A generalised linear model was used to test for differential expression across the following comparisons: (i) diagnosis specific effects (MDD vs CON; BPD vs CON; MDD vs BPD), (ii) transdiagnostic mood effects (MDD+BPD vs CON) and (iii) sex effects (male vs female, across all diagnoses). An interaction model (sex x diagnosis) was not examined due to limited power.

To assess the effect of potential confounding variables, a principal component analysis (PCA) was performed on the total gene expression profiles using all samples. PCA was conducted using demographic (age) and technical variables (RNA integrity number, brain pH, postmortem interval and RNA extraction batch). No strong clustering or separation patterns were observed for any of the variables across the first 4 principal components, indicating minimal contribution of these factors to overall variance (**Supplementary Figure S1**). Therefore, no covariates were included in the models to avoid overfitting given the small sample size. Statistical significance was determined using the Wald test and false discovery rate (p<0.05, log2FC<0 for downregulated genes and log2FC>0 for upregulated gene) was adjusted for multiple comparisons using the Benjamini-Hochberg correction.

### 2.6 Gene Enrichment Analysis

DEGs identified in the habenula were subjected to functional enrichment analysis using g:Profiler (40). A custom background gene list, defined as all genes detected in the RNA-seq dataset was applied. Overrepresentation analysis was performed using the Gene Ontology Biological Process (GO:BP), Gene Ontology Molecular Function (GO:MF) and Reactome databases. Complete enrichment results, including analyses across all databases available in g:Profiler, are provided in the supplementary materials. Correction for multiple comparisons was applied using the g:SCS algorithm, with significance set at an adjusted p value <0.05. Heatmap analyses were performed on variance-stabilising transformed (VST) counts and were generated for significant DEG (adjusted p-value < 0.05), with hierarchical clustering applied to genes. For presentation, pseudogenes and uncharacterised transcripts were removed.

### 2.7 Differential transcript usage (DTU) analysis

Differential transcript usage (DTU) was performed to identify changes in relative isoform expression that may occur independently of total gene expression changes. DTU can reveal functionally relevant isoform-level switching and usage that is not captured by differential gene expression analyses, and has been increasingly applied in transcriptomic studies of complex brain disorders (41,42)

DTU was assessed with IsoformSwitchAnalyzeR v2.8.0 (43) using DEXSeq (44). Only datasets with paired-end sequencing and a minimum sequencing depth of 30 million reads were included. Genes with low overall expression (mean TPM ≤ 1 in both mood and control groups), transcripts contributing less than 1% of total gene expression, and genes expressing a single transcript were excluded prior to testing. Given limited power for isoform-level analyses, DTU was restricted to the transdiagnostic mood (MDD+BPD vs control) and sex (male vs female) contrasts to maximise statistical sensitivity for detecting shared transcript-level alterations. Isoform switches were considered significant at a false discovery rate (FDR) < 0.05 and an absolute change in isoform fraction (|IF|) ≥ 0.1, corresponding to a minimum 10% change in transcript usage. Identified isoform switching genes were evaluated with CPAT (coding potential), PfamScan (protein domains), SignalP (signal peptides), and analyzeDeepTMHMM (transmembrane regions). Isoform switching events were defined as transcripts originating from the same gene showing opposing changes in usage of at least 10 % (|ΔIF|) ≥ 0.1 between mood and control or male and female samples. Significant isoform switches were classified as functionally relevant based on predicted differences in one or more molecular feature, including intron retention, coding potential, open reading frame structure, nonsense-mediated decay status, protein domain structure, or signal peptide presence.

## 3. Results

A PCA on the total gene expression profiles using all samples showed no clear clustering by diagnosis (**Supplementary Figure S2**). Notably, the PCA did reveal a strong separation of samples by sex (**Supplementary Figure S3**). One sample (BPD5) was isolated from the main distribution of cases, and the variance could not be explained by demographic or technical covariates. On this basis BPD5 was considered an outlier and excluded from the primary analysis (**Supplementary Figure S4**). Results from additional analyses including BPD5 are provided in the supplementary material.

### 3.1 Transcriptomic differences in the human habenula across mood disorders

DEG analysis identified changes in the habenula across both diagnostic and sex-based comparisons. Compared to controls, 60 genes (21 upregulated, 39 downregulated) were differentially expressed in MDD cases (FDR<0.05) (**Figure 2**). Comparison between BPD and control cases yielded a similar number, identifying 66 DEGs (41 upregulated, 25 downregulated BPD vs CON). Direct comparison between BPD and MDD cases identified only three differentially expressed transcripts, all of which were non-coding genes. These included two pseudogene-related loci (C22orf46P and PMS2CL) and one long intergenic non-coding RNA (LINC03104), none of which have established roles in habenula function or psychiatric neurobiology. The minimal differences observed between BPD and MDD are consistent with substantial overlap in depressive symptomatology and support a shared underlying molecular architecture across mood disorders. Consistent with the predefined transdiagnostic framework, analysis of the combined mood cohort (MDD+BPD) identified a substantially larger set of 378 differentially expressed genes (245 upregulated, 133 downregulated) relative to controls. More than 80% of genes identified as differentially expressed in the individual diagnostic comparisons were recapitulated in the combined mood analysis, indicating that the majority of diagnosis-specific signals converge within a shared molecular signature across mood disorders.

**Figure 2.**
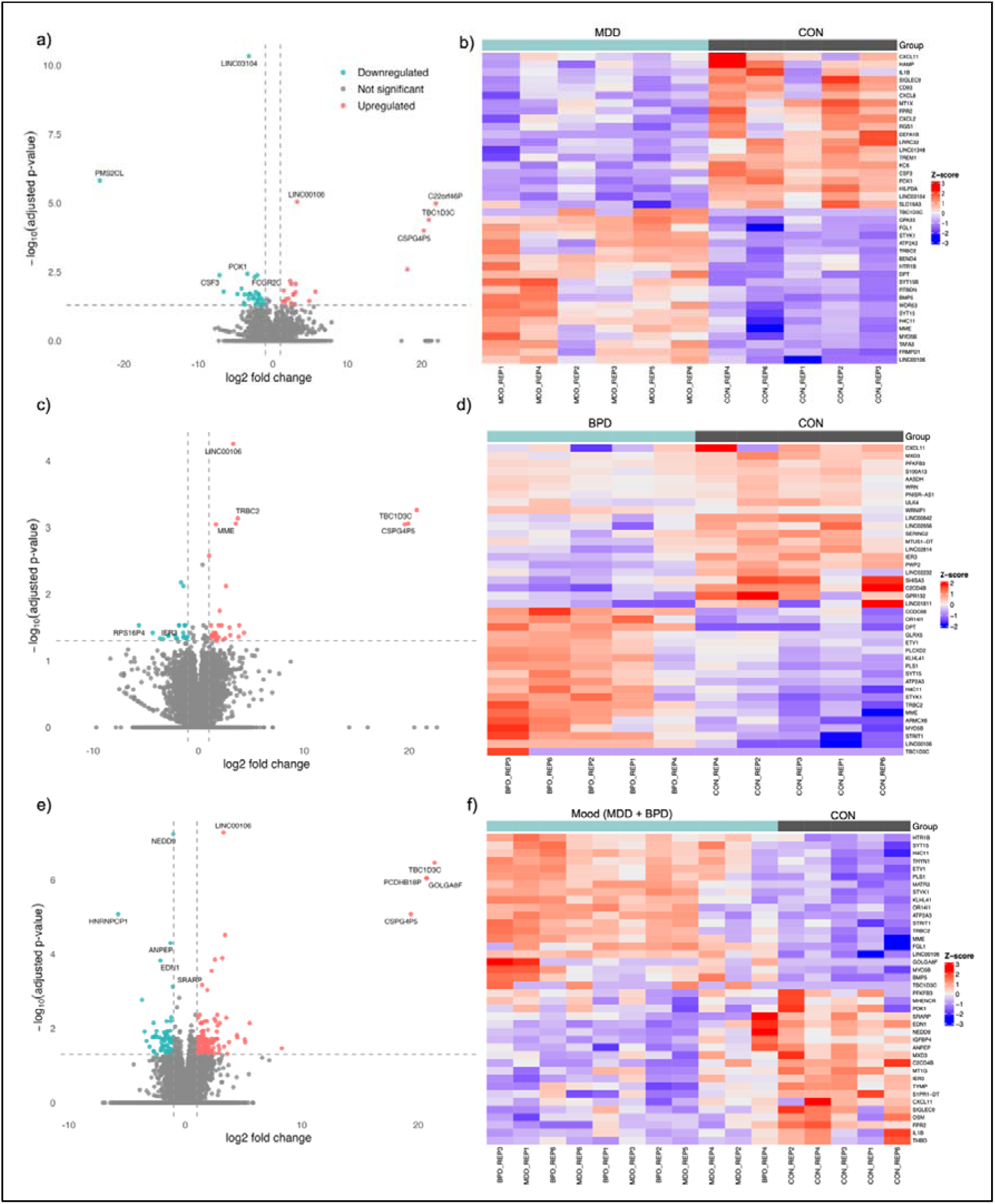
Visualisation of differential expressed genes in MDD versus CON (a-b) BPD versus CON (c-d) and Mood (MDD + BPD) versus CON (e-f). The left panel depicts volcano plots of differentially expressed genes with significantly upregulated (red) and downregulated (blue) genes highlighted (padj <0.05,|log2FC| ≥ 1)). The right panel shows heatmaps of row- z-scaled log₂(TPM+1) expression values for the top 20 upregulated and top 20 downregulated differentially expressed genes. Warmer colours indicate higher relative expression, and cooler colours indicate lower relative expression.

Given the strong convergence of differential expression across mood cases, downstream functional analyses focused on the transdiagnostic DEG set. Functional enrichment analysis of DEGs identified biological processes consistent with altered habenula signalling in mood disorders. Genes upregulated in the mood cohort (MDD+BPD vs CON) were significantly enriched for pathways related to excitatory neurotransmission, including potassium channel activity, Wnt signalling, calcium ion homeostasis and calmodulin binding indicating enhanced ion-dependent signalling processes (**Figure 3**). In parallel, enrichment of GABAergic signalling pathways was observed, largely driven by higher expression of *GABBR2* (p<0.05). Notably, *GABAB2* receptors in the habenula have previously been reported to localise predominantly to presynaptic GABAergic terminals, where their activation reduces GABA release, a mechanism that may functionally disinhibit excitatory output (45). Conversely, downregulated genes were enriched for processes related to metal ion binding and homeostasis, a finding not previously identified in preclinical habenula models. Disruption of metal ion regulation has, however, been implicated in broader psychiatric and neurobiological contexts (46).

**Figure 3.**
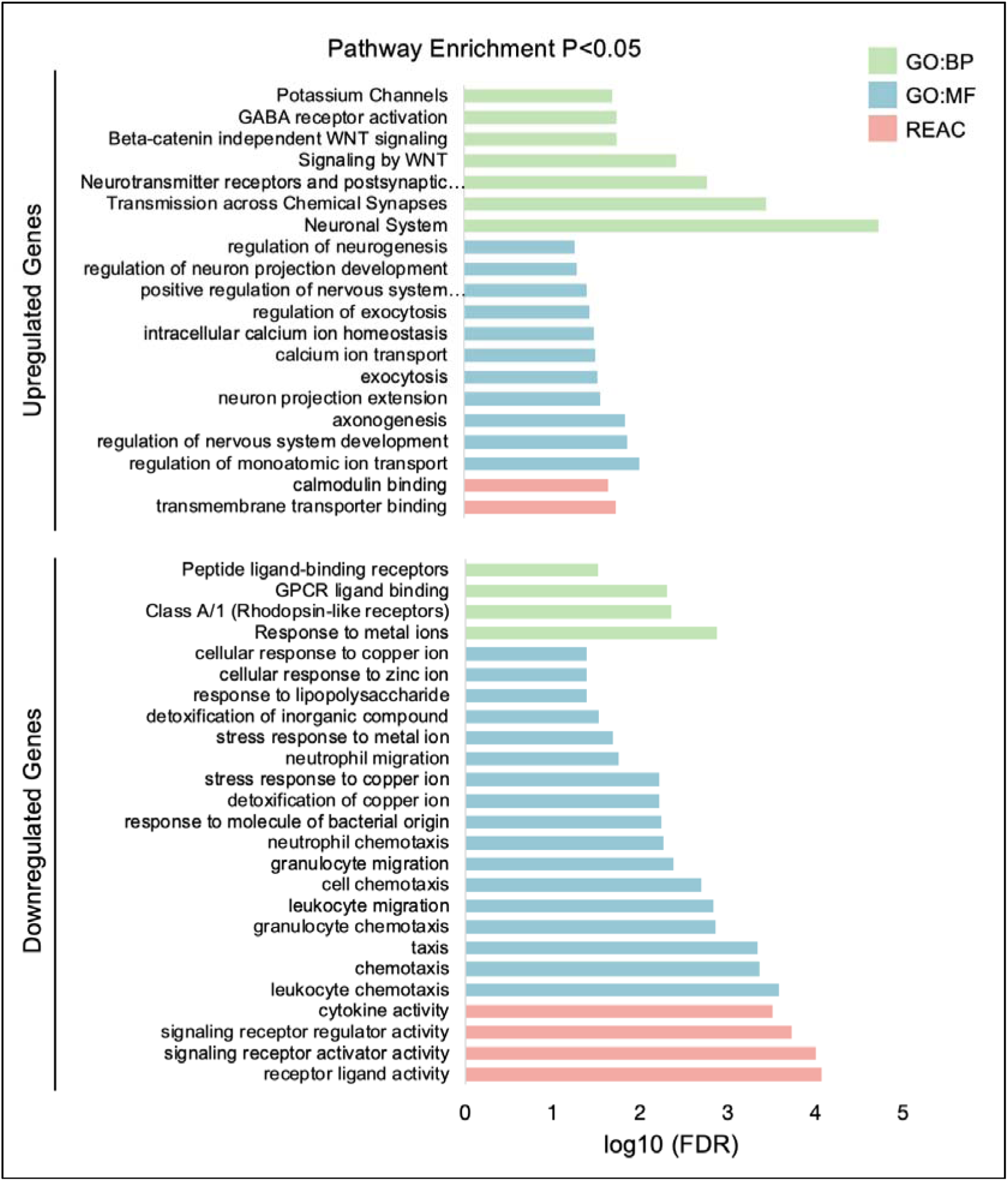
Functional enrichment of upregulated and downregulated genes in mood (MDD + BPD) versus control cases. Overrepresentation analysis was performed in g:Profiler using Gene Ontology Biological Process (GO:BP), Gene Ontology Molecular Function (GO:MF), and Reactome (REAC) databases. Enriched terms for upregulated genes (top) and downregulated genes (bottom) are shown, with bars representing significantly enriched pathways (adjusted *p* < 0.05, g:SCS correction) and coloured by database.

Differential transcript usage (DTU) analysis identified 49 significant isoform switches across 41 genes in the human habenula (Supplementary Table 13). Isoform switching refers to a change in the relative abundance of transcript variants without corresponding changes in total gene expression. Only two DTU genes overlapped with those identified by differential gene expression (DEG) analysis, while the remaining genes exhibited isoform switching in the absence of detectable changes at the gene-expression level. Among these DTU-affected genes, *MGLL,* an enzyme central to endocannabinoid signalling, exhibited significant isoform switching, with increased usage of ENST00000265052 in mood cases. Isoform-specific changes were also observed in the *KCNKJ10*, a potassium channel transcript, strongly implicated in habenula excitability, with increased usage of ENST00000340700. Similarly, *CAMK2B* and *RBFOX3* (NeuN), genes with known neuronal functions, showed significant isoform switching despite unchanged total gene expression (**Figure 4**). Specifically, *CAMK2B* isoform ENST00000700289 demonstrated increased usage in mood cases (p < 0.05). Structural annotation indicates that this transcript modifies the coding sequence relative to the canonical isoform, potentially modifying CaMKIIβ kinase domain architecture. Likewise, although total *RBFOX3* expression remained stable, ENST00000580115 exhibited increased relative usage in mood samples. Given that *RBFOX3* encodes a neuron-specific RNA-binding protein containing an RNA Recognition Motif (RRM) essential for splice-site recognition, altered isoform usage may modify protein domain composition and consequently influence downstream splicing regulation. Together, these findings indicate that isoform-level regulatory shifts represent a core and transdiagnostically conserved feature of molecular alterations in the human habenula across mood cases. Rather than altering the overall abundance of key neuronal genes, mood related pathology may preferentially reshape structural variants of proteins involved in E:I balance, revealing a layer of molecular dysregulation not captured by conventional DEG analysis. A complete list of significant DTU events between mood and control groups is provided in the supplementary material.

**Figure 4.**
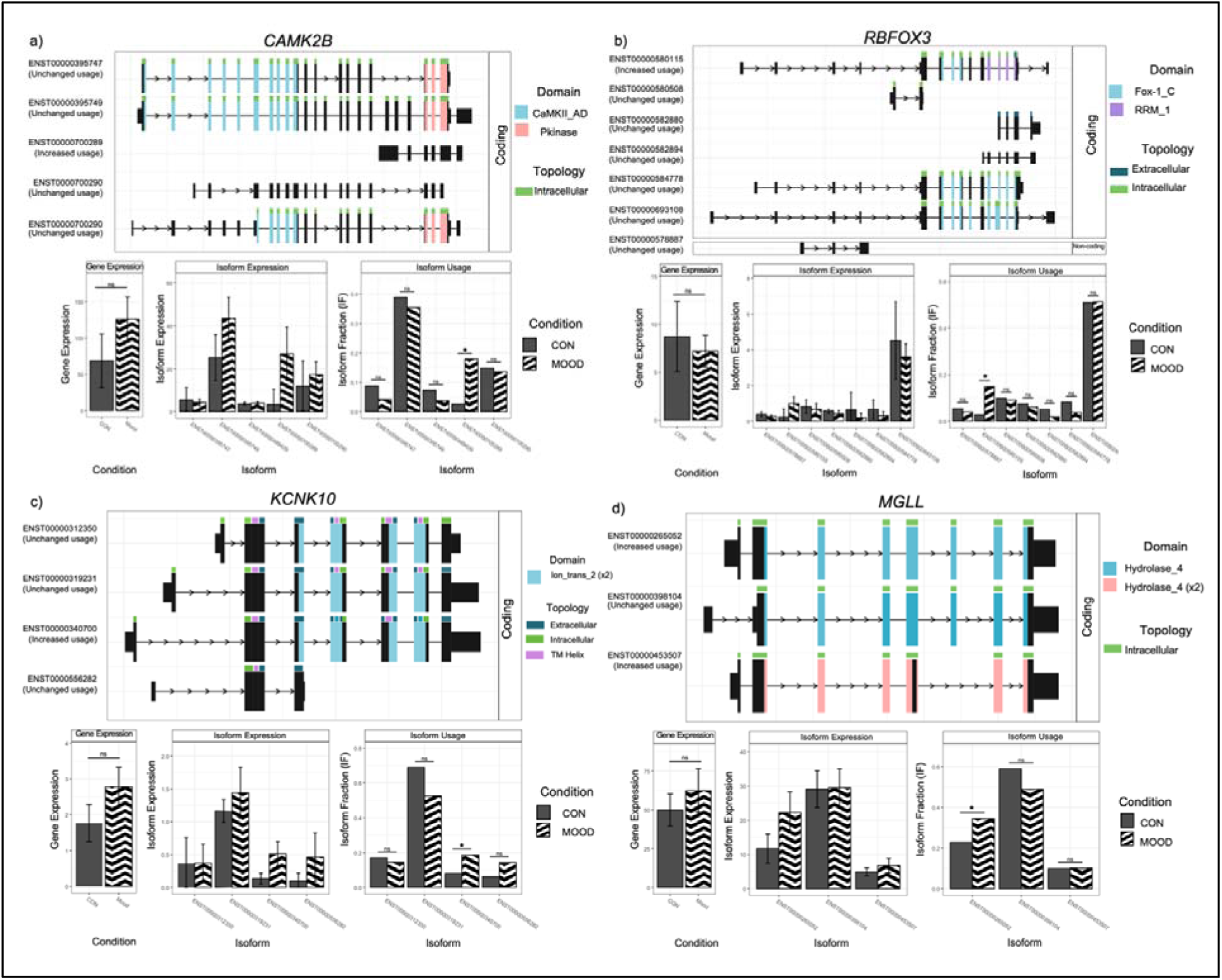
Isoform switch plots illustrating differential transcript usage in the human habenula in control vs mood disorder cases. Differential transcript usage (DTU) analysis revealed isoform-specific alterations in *CAMK2B* (a), *RBFOX3* (b), *KCNK10* (c), and *MGLL* (d) in post-mortem human habenula tissue from mood (BPD + MDD) compared with controls. Transcript schematics depict exon–intron organisation, predicted protein domains, and membrane topology (top), with corresponding bar plots showing total gene expression, isoform-level expression, and isoform fraction (IF) for each condition (bottom). Bars represent mean with 95% confidence intervals. *: *p* < 0.05; ns, not significant.

### 3.2 Transcriptomic sex differences in the human habenula

Additional sex stratified analysis identified 67 DEGs (12 downregulated, 55 upregulated in males compared to females) (**Figure 5**). Of these, 48 genes were located on either the x or y sex chromosomes, while 19 were autosomal sex differences. Canonical sex chromosome markers showed the expected expression patterns, with female biased expression of *XIST*, *JPX*, *KDM5C*, and *DDX3X*, and male biased expression of *UTY*, *TBL1Y*, *ZFY*, and *DDX3Y*, providing internal validation of the differential expression analysis. Beyond these markers, several autosomal genes exhibited sex dependent expression, indicating broader sexual dimorphism in habenula gene regulation. Notably, *GRP151*, an inhibitory g-protein coupled receptor highly enriched in habenula neurons (33), was expressed at higher levels in females, whereas *PDE8B*, a negative regulator of cAMP signalling, was higher in males.

**Figure 5.**
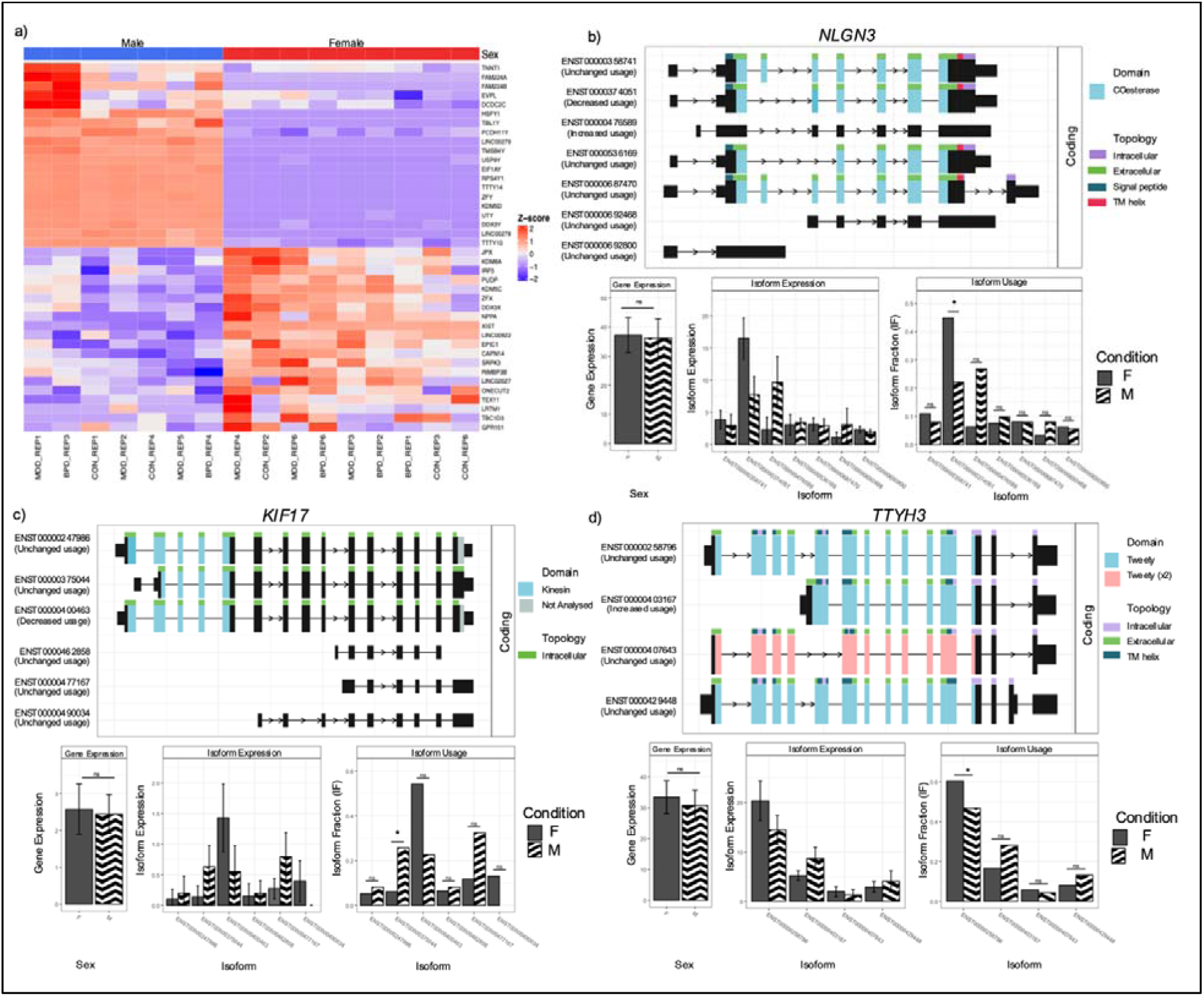
Differential gene expression and differential transcript usage in the human habenula in females vs males cases. **(a)** Heatmap of row-scaled log₂(TPM+1) expression values for the top 20 upregulated and top 20 downregulated differentially expressed genes in male versus female cases. Warmer colours indicate higher relative expression and cooler colours indicate lower relative expression.Differential transcript usage (DTU) analysis revealed isoform-specific alterations in *NLGN3* **(b)**, *KIF17* **(c)**, and *TTYH3* **(d)** in post-mortem human habenula tissue from male vs female cases. Transcript schematics depict exon–intron organisation, predicted protein domains, and membrane topology (top), with corresponding bar plots showing total gene expression, isoform-level expression, and isoform fraction (IF) for each condition (bottom). Bars represent mean with 95% confidence intervals. * *p* < 0.05; ns, not significant.

DTU analysis identified 18 significant isoform switches between males and females, all of which were independent of total gene expression changes (Supplementary Table 14). Notably, although total *NLGN3* expression did not differ between sexes, females exhibited increased usage of ENST00000374051, a transcript predicted to lack canonical cholinesterase-like domain features and key membrane topology elements, including signal peptide and transmembrane regions. This redistribution toward a structurally truncated isoform suggests a potential shift in synaptic protein composition despite unchanged total gene levels. Given that *NLGN3* functions as a postsynaptic adhesion molecule critical for synapse formation and maintenance, altered isoform usage may alter protein localisation, membrane integration, or synaptic adhesion properties despite stable overall gene expression. In contrast, males exhibited increased usage of ENST00000400463 of *KIF17,* a kinesin motor protein involved in NMDA receptor trafficking, suggesting sex-dependent modulation of post-synaptic receptor transport (47) (**Figure 4**). A complete list of significant DTU events between male and females is provided in the supplementary material.

## 4. Discussion

Understanding the molecular landscape of the human brain is essential for determining how region-specific alterations contribute to circuit and system level dysfunction in mood disorders. While preclinical research has provided critical insights into the habenula’s role in the neurobiology of depression, aspects of its molecular architecture are not conserved across species, highlighting the need for direct investigation of the human habenula in psychiatric illness. Here, we present the first transcriptomic profiling of the human habenula across mood disorders, identifying a dominant transdiagnostic molecular signature shared between MDD and BPD, alongside convergence with pathways previously implicated in preclinical models of depression and the identification of novel transcriptional changes. Importantly, our findings also reveal biological sex as a major source of transcriptomic variation in the human habenula, extending beyond expected sex-chromosome–linked effects.

### 4.1 Transcriptomic alterations in the human habenula in mood disorders

Consistent with the predefined transdiagnostic framework, the dominant molecular signal in the human habenula was shared across MDD and BPD, with enrichment of pathways related to excitatory neurotransmission, including potassium channel activity, calcium ion transport and calmodulin binding. Preclinical evidence implicates potassium channel dysfunction as a central driver of elevated habenula neuronal excitability (48). The inward rectifying potassium channel, Kir4.1 encoded by *KCNJ10*, expressed on astrocytes, regulates the degree of membrane polarisation and extracellular potassium buffering (19). In mice, overexpression of Kir4.1 in the habenula is shown to induce neuronal bursting and subsequent depressive-like phenotypes that are reversed by Kir4.1 ablation or inhibition (19). Notably, preclinical models have identified Kir4.1 inhibitors as novel rapid-acting antidepressant candidates with favourable safety profiles, supporting astrocytic potassium channels as a promising therapeutic target (49,50). In the present study, *KCNJ10* exhibited significant isoform switching in mood disorder cases, suggesting altered astrocyte-mediated potassium regulation at the transcript level in the human habenula. Consistent with this, additional potassium channel genes were elevated, including the neuronally expressed inwardly rectifying channel *KCNJ12* and several voltage-gated subtypes (*KCNQ3, KCNH5, KCNF1*), suggesting broader disruption of markers associated with intrinsic potassium signalling across cell types.

Beyond potassium-mediated regulation of excitability, preclinical studies have consistently implicated glutamatergic and calcium-dependent signalling as an additional mechanism underlying habenula hyperactivity (14,18). Elevation of CaMKIIβ, which promotes glutamatergic signalling, in the LHb is sufficient to induce depressive-like behaviours in rodents and is required for their maintenance. In contrast, blockade or pharmacological downregulation (via ketamine) of CaMKIIβ normalises LHb firing and reverses behavioural despair in rodent models (14,51). In the present study, relatively few genes encoding glutamatergic receptors were differentially expressed between mood disorder and control cases. However, genes related to calcium ion transport and homeostasis were broadly elevated, indicating altered intracellular signalling rather than changes in postsynaptic glutamate receptor expression. Consistent with this, although *CAMK2B*, the gene that encodes for CaMKIIβ showed no detectable differences in total gene expression, differential transcript usage revealed isoform-specific regulation, including increased usage and switching of specific *CAMK2B* variants in mood disorder cases. Different *CAMK2B* isoforms encode distinct functional domains that shape protein-protein interactions and signalling, allowing neuronal function to change without alterations to overall gene expression (52,53). Given that CaMKIIβ is enriched in glutamatergic neurons and functions as a key regulator of calcium signalling, synaptic plasticity and excitatory transmission (51), these findings suggest that glutamatergic dysregulation in the human habenula may be mediated through transcript-level regulation of intracellular signalling components rather than widespread changes in glutamate receptor expression. Together, these findings suggest that alterations in intrinsic excitability and intracellular signalling represent convergent molecular mechanisms across mood disorders within the human habenula. While increased sample size contributes to statistical sensitivity in the combined analysis, the high degree of overlap between individual diagnostic and transdiagnostic comparisons supports a biologically meaningful shared signal rather than a purely power-driven effect.

In parallel, *MGLL*, the principal enzyme responsible for degradation of the endocannabinoid 2-arachidonoylglycerol (2-AG), exhibited significant isoform switching in mood disorder cases. In cortical and hippocampal regions, increased 2-AG signalling enhances CB1 receptor–mediated retrograde inhibition of glutamate release, thereby dampening neuronal excitability (54,55). In contrast, within the lateral habenula, 2-AG signalling has been shown to paradoxically potentiate neuronal firing, likely through preferential CB1-mediated suppression of inhibitory GABAergic neurons, resulting in net disinhibition (55). Given that the functional consequences of endocannabinoid signalling depend on the cellular and synaptic localisation of CB1 receptors, altered isoform-level regulation of *MGLL* in the human habenula may shift the balance between inhibitory and excitatory signalling in a manner distinct from other brain regions. Although isoform switching does not directly equate to altered enzymatic activity, these exploratory findings underscore the need for region-specific investigation of endocannabinoid regulation in mood disorders, particularly in the context of the habenula, which has been reported to exhibit hyperactivity rather than the hypoactivity more typically observed in cortical regions (56,57).

Genes upregulated in mood disorder cases were enriched for Wnt and β-catenin–dependent signalling pathways. Preclinical developmental studies demonstrate that precise temporal regulation of Wnt activity is essential for normal habenular neuron differentiation; however, how perturbations of this pathway influence habenula function later in life, or whether Wnt signalling is dysregulated in the mature habenula across psychiatric conditions, remains unclear (58,59). Notably, preclinical studies show that stress alters the expression of Wnt pathway components and that several Wnt-related molecules are responsive to antidepressant treatment, implicating this pathway in depression-related behaviour (60–62). Wnt signalling also converges on intracellular cascades such as MAPK/ERK and PI3K, which are disrupted in both postmortem human (63) and preclinical depression studies (64), although the extent to which these changes reflect habenula specific mechanisms versus broader system level alterations is unclear.

Conversely, downregulated genes were enriched for processes related to metal ion binding and homeostasis, a pathway not previously highlighted in preclinical models of habenula dysfunction. Although the role of metal and trace element regulation in the habenula remains poorly characterised, disruption of these processes has been linked more broadly to neuropsychiatric phenotypes through mechanisms involving oxidative stress, mitochondrial dysfunction and impaired ATP synthesis (65–67). Consistent with this, clinical studies have associated zinc deficiency with MDD (67,68) and copper dysregulation with BPD and schizophrenia (46).

At glutamatergic synapses, zinc is a key modulator of excitatory transmission, exerting inhibitory effects on GluN2A-containing NMDA receptors (69) and influencing AMPA receptor function (70). Zinc availability is tightly regulated by transporters that control vesicular loading and intracellular distribution. Reduced expression of the vesicular zinc transporter *ZnT3* has previously been reported in the prefrontal cortex of individuals with depression and is thought to reflect diminished synaptic zinc signalling (71). While *ZnT3* expression was unchanged in the present study, we observed reduced expression of the zinc importer *ZIP10*, alongside significant downregulation of multiple metallothioneins (*MT1A, MT1X, MT2A, MT3*), which act as critical intracellular zinc- and copper-binding proteins. Notably, previous transcriptomic analyses in bipolar disorder have identified metallothionein and metal ion binding pathways among the most significantly enriched gene categories, driven in part by *MT1X and MT2A* (72). However, this study reported increased expression of metallothionein-related genes in the PFC, with no corresponding changes observed in the hippocampus (72). In contrast, our findings demonstrate reduced expression within the habenula. This divergence likely reflects region-specific functional organisation, particularly given the opposing roles of the PFC and habenula in mood regulation, and suggests that dysregulation of metal ion homeostasis may contribute more broadly to circuit-level dysfunction. Given the central role of metallothioneins in buffering intracellular metal ions and limiting oxidative stress, their coordinated reduction with altered zinc transporter expression supports disruption of intracellular metal ion homeostasis in the human habenula in mood disorders. Together, these findings identify metal ion homeostasis as a novel and underexplored molecular pathway of the human habenula; however, whether these changes contribute directly to mood disorder pathology or arise secondary to broader alterations in habenula activity is unclear.

### 4.2 Sex-biased transcriptional regulation in the human habenula

Sex differences are increasingly recognised as fundamental to the neurobiology of psychiatric disorders, yet the molecular mechanisms through which biological sex shapes vulnerability, symptom expression and treatment response remain poorly defined. In the present study, principal component analysis revealed no clear separation by diagnosis; however, samples clustered strongly by sex, indicating biological sex as a major source of transcriptomic variation in the habenula. Beyond expected sex-chromosome effects, several autosomal genes with functional relevance to habenula activity showed sex-biased expression. Notably, *GPR151*, a Gi/o-coupled inhibitory G-protein–coupled receptor selectively enriched in lateral habenula neurons (33), was expressed at higher levels in females. Preclinical studies demonstrate that GPR151 couples to Gαo to suppress cAMP signalling, modulates synaptic transmission and plasticity, and critically regulates habenula excitability (73). Accordingly, increased *GPR151* expression in females may reflect sex-specific regulation of presynaptic inhibitory control within LHb circuits, although whether this represents a protective or compensatory response remains unclear. Females also showed increased usage of the *TTYH3 isoform* ENST00000403167, a calcium-activated chloride channel implicated in ionic stability and the modulation of neuronal excitability (74). Whereas males exhibited higher usage of specific isoforms of *KIF17*, involved in NMDA receptor trafficking (47). Collectively, these transcriptional and isoform-level differences are consistent with sex-specific modulation of excitatory/inhibitory balance within habenula neurons. Whether these patterns reflect primary biological differences, compensatory adaptations, or clinically relevant mechanisms remains unclear.

### 4.3 Limitations and future directions

The present study is limited by the small sample size which reduced statistical power and precluded formal testing of sex x diagnosis interaction effects, analyses were therefore restricted to main effects of diagnosis and sex. However, the sample size is comparable to other postmortem human brain RNA-sequencing studies in psychiatry (75,76) and consistent with prior postmortem investigations of the human habenula (32). Importantly, achieving large, sex balanced cohorts sufficient to interrogate interaction effects remains challenging. The substantial financial costs associated with high-throughput transcriptomic methodologies, coupled with availability of postmortem human brain tissue constrains sample sizes. In addition, the analytical and computational demand increases considerably when modelling stratified effects across both diagnosis and sex. Despite these challenges, integrating sex as a biological variable is essential for advancing mechanistic insight into psychiatric disorders

Although the cohort included cases with major depressive disorder and bipolar disorder, the bipolar cohort was predominantly characterised by depressive symptomology with limited history of mania. Consequently, our findings may reflect molecular processes associated with depressive symptomatology across mood disorders, aligning with a transdiagnostic framework, in which shared features reflect common underlying biology rather than distinct diagnostic categories. Future studies incorporating a larger cohort are crucial to resolve molecular nuances between diagnostic groups.

In addition, transcriptomic analyses of postmortem tissue capture a late stage of disease, and it is unclear whether the observed molecular changes represent primary pathological drivers of depression or secondary downstream effects. All mood disorder cases had a documented history of antidepressant exposure, and while medication effects on habenula molecular architecture remain poorly characterised, their potential contribution to the observed transcriptional changes cannot be excluded. Future studies incorporating medication-naïve cohorts or stratified analyses will be important to elucidate possible treatment-related effects. In addition, alterations in the habenula have previously been reported in MDD cases who died by suicide compared to MDD cases who did not, suggesting clinical heterogeneity, including suicide status, may influence molecular changes observed in the region (77).

Furthermore, the use of bulk RNA sequencing does not allow for identification of the specific cell types that may be contributing to the observed transcriptional changes. Although tissue was highly enriched for the habenula, some contamination from adjacent thalamic regions is expected and the small size of the structure prevented differentiation between the medial and lateral subdivisions. Similarly, while emerging evidence suggests functional and molecular lateralisation within the habenula, hemispheres were homogenised in the present study to maximise RNA yield, potentially obscuring hemisphere-specific effects (78,79).

Finally, transcriptomic alterations do not necessarily translate to protein level changes or functional outcomes, highlighting the need for future studies integrating proteomics, single cell approaches and molecular imaging to elucidate the biological significance of these molecular changes. Furthermore, validation in an independent cohort or through complementary molecular approaches was not feasible due to the limited availability of human habenula tissue. Consequently, these finding should be interpreted as preliminary, providing a valuable framework for future studies aimed at functionally characterising the molecular pathways identified.

### 4.4 Conclusions

This study provides the first transcriptomic characterisation, to our knowledge, of the postmortem human habenula in mood disorders. Notably, the predominant signal identified was transdiagnostic, with shared molecular alterations observed across major depressive disorder and bipolar disorder. These findings provide preliminary evidence that molecular pathways implicated in preclinical models of depression are conserved in the human brain while also shedding light on novel transcriptomic and sex specific alterations that had previously not been described. Alterations in genes associated with ion channel signalling, calcium-dependent excitability and astrocyte-mediated potassium regulation support dysregulated habenula functioning and hyperexcitability in the neurobiology of depression. Notably, the identification of genes associated with disrupted metal ion homeostasis and intracellular trace metal handling highlights a previously unrecognised molecular pathway in the human habenula, extending current models beyond major neurotransmitter systems. In parallel, widespread robust sex-dependent differences in gene expression and isoform usage identify sex as a fundamental determinant of habenula molecular organisation, with possible implications for vulnerability and treatment response. The identification of isoform specific regulation in key signalling pathways further underscores the value of transcript level analyses for capturing biologically relevant mechanisms not evident from gene expression alone. Together, these findings advance our understanding of habenula dysfunction in mood disorders and identify candidate molecular pathways to inform future mechanistic investigations and the development of more targeted, sex-informed therapeutic approaches.

## Acknowledgements

Human tissue was provided by The Netherlands Brain Bank (Netherlands Institute of Neuroscience, Amsterdam). RNA sequencing was performed by the Ramaciotti Centre of Genomics (University of New South Wales, Sydney). This research was undertaken with the assistance of resources and services from the National Computational Infrastructure (NCI), which is supported by the Australian Government, including access to the Gadi high-performance computing system. Sarah Cameron and Marnie L Maddock were supported by an Australian Government Research Training Program (RTP) Scholarship. KAN is funded by the NSW Ministry of Health, Office of Health and Medical Research.

## Author Contributions

SC, KN and KWG conceptulised and designed the study. SC, MM and SW developed the methodology. SC and MM performed the statisical analysis. SC and KN prepared the original draft of the manuscript. All authors (SC, MM, SW, KWG and KN) reviewed and approved the final version of the manuscript.

## Data Availability

Additional DEG, DTU, and gene enrichment data is provided in the supplementary material. All code developed for analysing the data is available at: https://github.com/smc331/habenula-dtu; https://github.com/smc331/habenula-DEG.

