## Supplementary Figures for "Transcriptomic profiling of the human habenula reveals a shared molecular architecture across mood disorders"

**Table of Contents**

**Supplementary Figure S1.** PCA of gene expression in enriched human habenula labelled by demographic and technical variables.

**Supplementary Figure S2.** PCA of gene expression in enriched human habenula samples labelled by diagnosis.

**Supplementary Figure S3*.*** **PCA of gene expression in enriched human habenula samples labelled by biological sex.**

**Supplementary Figure S4.** **PCA of gene expression in enriched human habenula samples before (a) and after (b) outlier removal.**

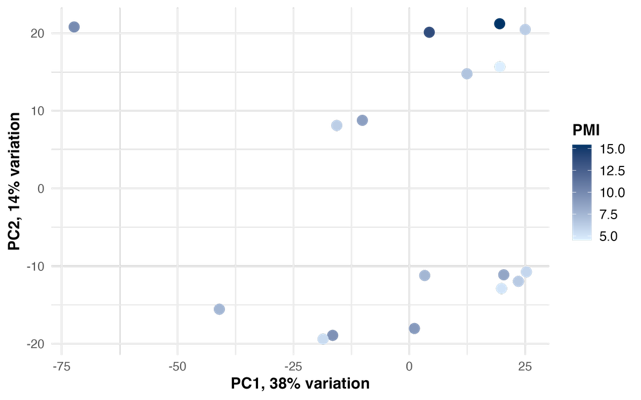

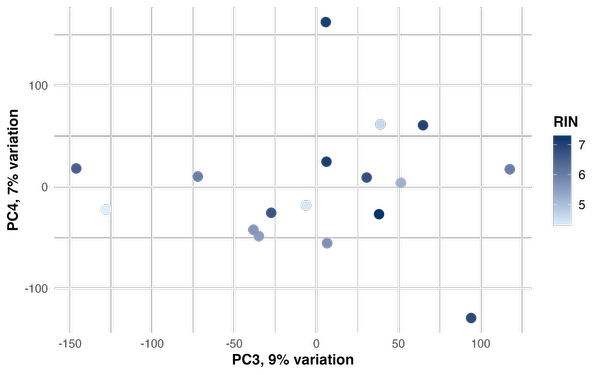

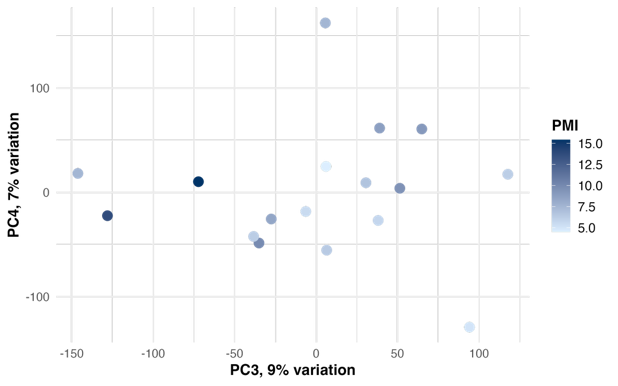

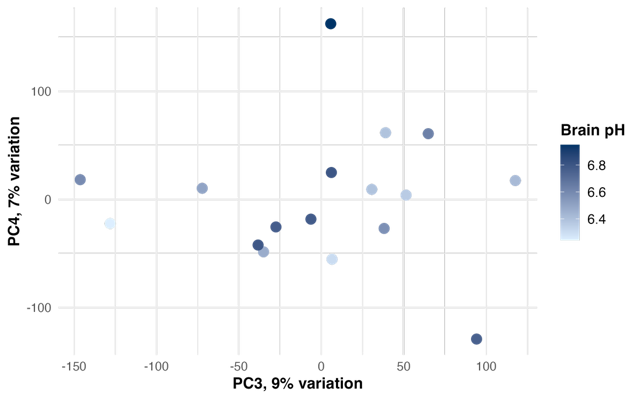

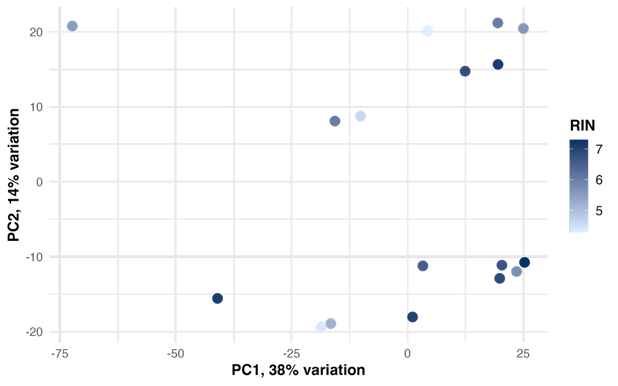

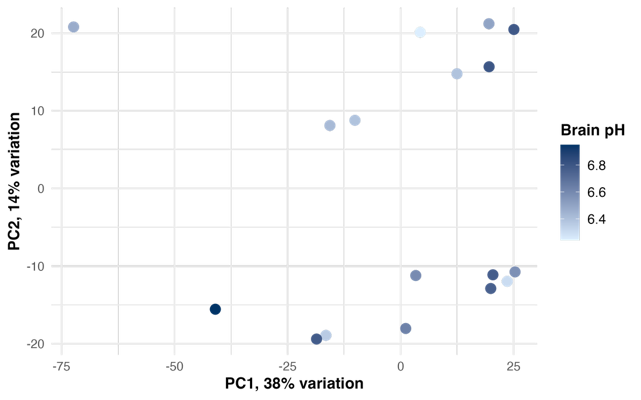

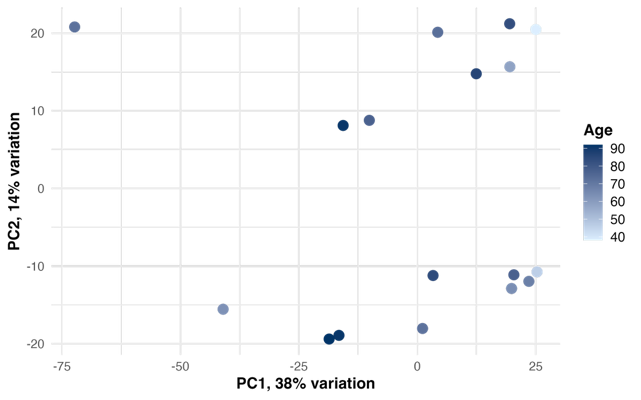

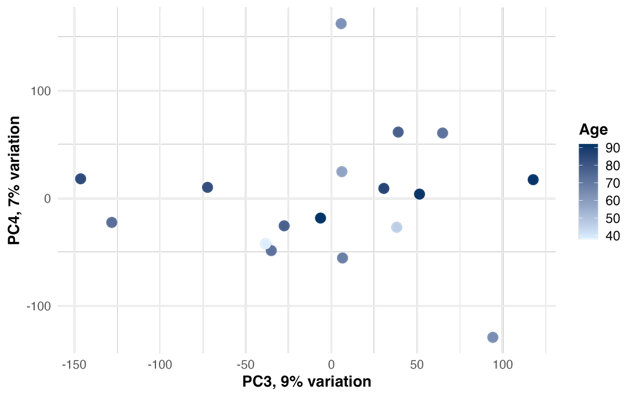

**Supplementary Figure S1*.*** PCA of gene expression in enriched human habenula labelled by demographic and technical variables. Scatterplots show PC1 versus PC2 and PC3 versus PC4, with samples coloured by age (a-b), brain pH (c-d), postmortem interval (PMI (e-f) and RNA integrity number (RIN) (g-h). No systematic clustering or consistent gradients were observed across the first four principal components for any variable.

h)

g)

e)

c)

a)

f)

d)

b)

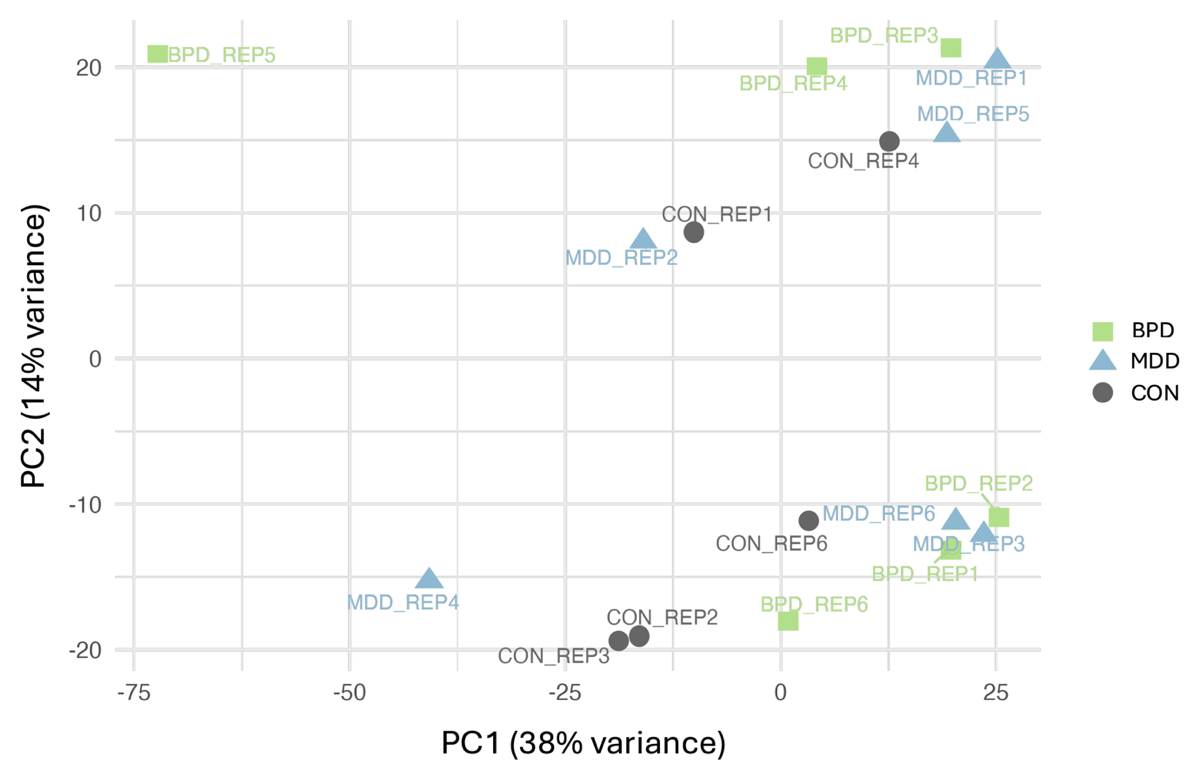

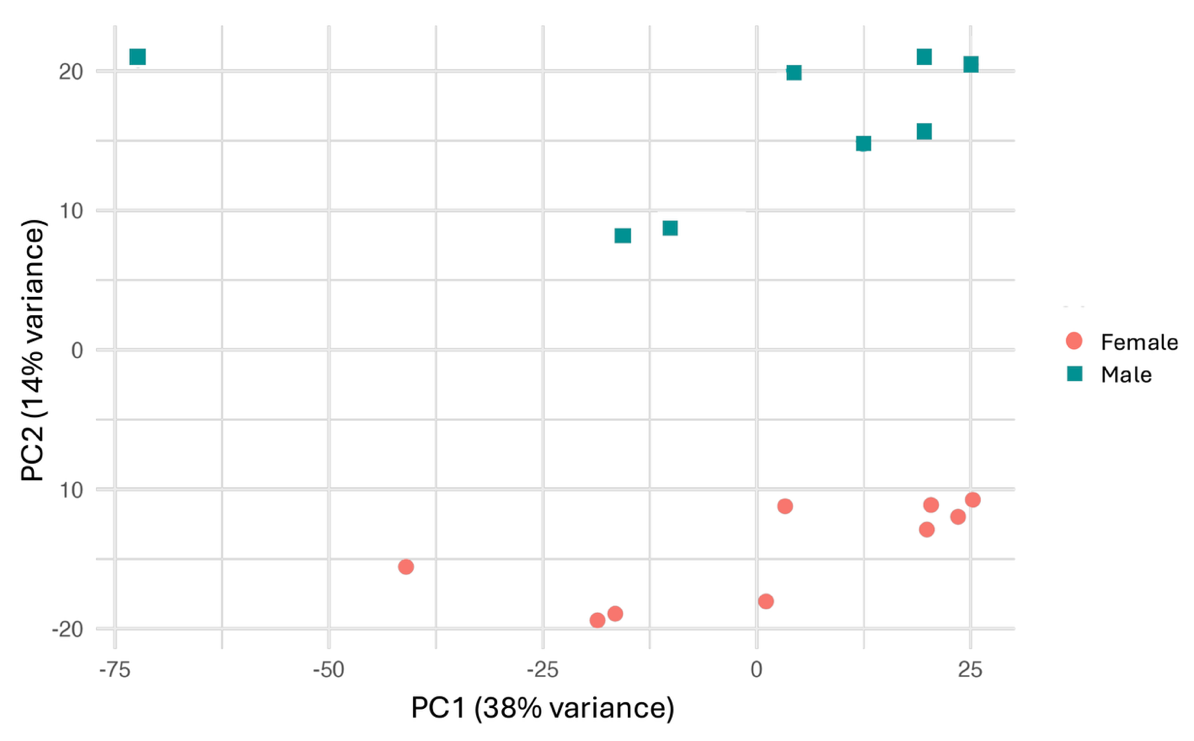

**Supplementary Figure S2*.*** **PCA of gene expression in enriched human habenula samples labelled by diagnosis.** Colours and shapes denote diagnostic groups (green/square = BPD, blue/triangle = MDD, grey/circle = CON).

**Supplementary Figure S3*.*** **PCA of gene expression in enriched human habenula samples labelled by biological sex.** Colours and shapes denote sex (blue/square = male, orange/circle = female).

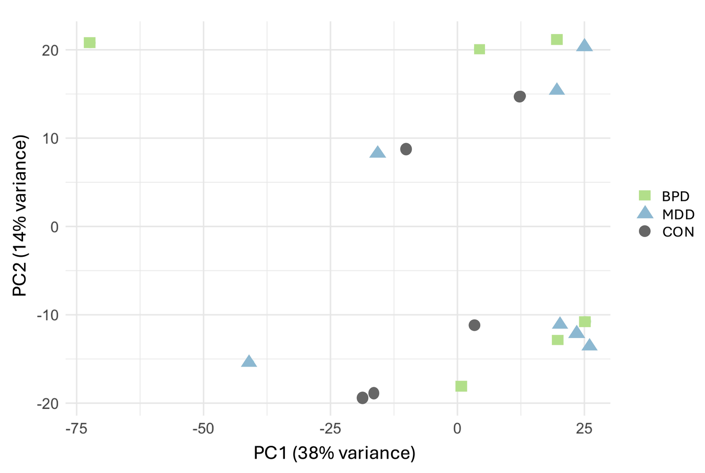

a)

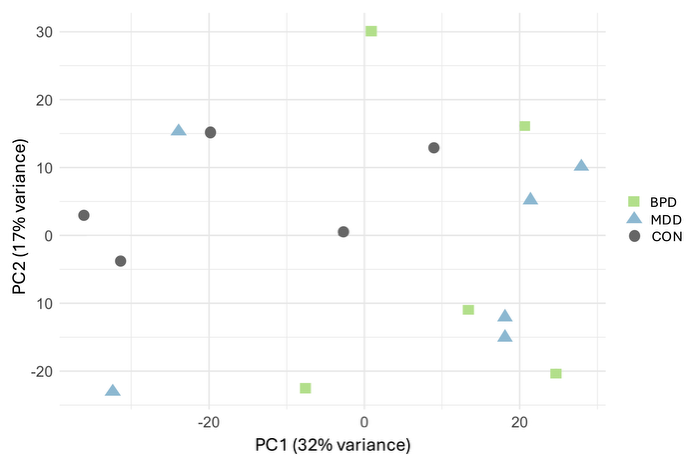

**Supplementary Figure S4.** **PCA of gene expression in enriched human habenula samples before (a) and after (b) outlier removal.** Colours and shapes denote diagnostic groups (green/square = BPD, blue/triangle = MDD, grey/circle = CON).

b)
